# A Novel Machine Learning Method for Mutational Analysis to Identifying Driver Genes in Breast Cancer

**DOI:** 10.1101/2022.11.20.517205

**Authors:** Golnaz Taheri, Mahnaz Habibi

## Abstract

Breast cancer has emerged as a severe public health issue and one of the main reasons for cancer-related mortality in women worldwide. Although the definitive reason for breast cancer is unknown, many genes and mutations in these genes associated with breast cancer have been identified using developed methods. The recurrence of a mutation in patients is a highly used feature for finding driver mutations. However, for various reasons, some mutations are more likely to arise than others. Sequencing analysis has demonstrated that cancer-driver genes perform across complicated pathways and networks, with mutations often arising in a modular pattern. In this work, we proposed a novel machine-learning method to study the functionality of genes in the networks derived from mutation associations, gene-gene interactions, and graph clustering for breast cancer analysis. These networks have revealed essential biological elements in the vital pathways, notably those that undergo low-frequency mutations. The statistical power of the clinical study is considerably increased when evaluating the network rather than just the effects of a single gene. The proposed method discovered key driver genes with various mutation frequencies. We investigated the function of the potential driver genes and related pathways. By presenting lower-frequency genes, we recognized breast cancer-related pathways that are less studied. In addition, we suggested a novel Monte Carlo-based algorithm to identify driver modules in breast cancer. We demonstrated our proposed modules’ importance and role in critical signaling pathways in breast cancer, and this evaluation for breast cancer-related driver modules gave us an inclusive insight into breast cancer development.

## 1 Introduction

Breast cancer is a considerably common type of cancer in women worldwide [1], and it can be inherited, which can be seen as a hazard killer in aggregated families. Some high and low-frequency mutation combinations drive this inheritance, such as BRCA1, BRCA2, P53, PALB2, CHEK2, and ATM, which confer an elevated lifetime risk of breast cancer. The common mutations at more than seventy loci were identified through large-scale replication studies [2]. Breast cancer has been eradicated only through tumor surgery and chemotherapy. At the same time, surgery is the only corrective approach for localized tumors, revealing a high risk of advanced metastasis [3]. Because of these limitations, new methods have been trying to develop breast cancer’s primary and secondary prevention strategies. On the other hand, women with BRCA1/2 germline mutations or a history of breast cancer are the only ones who can get a bilateral mastectomy [4, 5]. The high and low mutated genes have been used as the target in numerous chemoprevention techniques, including the use of selective estrogen receptor modulators, micronutrients, and anti-estrogen drugs [2].

The global approaches for analyzing networks try to understand and predict mechanisms in biological phenomena to investigate complex interactions in these biological networks. These approaches examine how various essential functions are exhibited outside of the association among their components. Large-scale omics data, high throughput gene expression data, and efficient integrative methods that have been developed are increasingly being used to map genes associated with specific biological functions or diseases [6]. Mutation in cancer plays an essential role in driving the associated genes to force ultimately defective protein translation, which in turn disturbs normal cell functioning. These genes are commonly used as target genes [7]. It has been reported that the breast cancer network has more than 6000 genes, which are involved in 7532 biological processes and 1930 molecular functions [8]. As a result, it is challenging to understand the network’s complex structure [9]. The emergent modular nature of the currently studied networks, including the protein-protein interaction (PPI) network, allows us to use bottom-up approach methods to investigate the different components, such as sub-networks, modules, and their associations at different levels. These network modules enable us to discover the underlying principle of breast cancer progression and bring the idea of searching for essential target genes into the well-organized breast cancer network [10]. Modules can be recursively divided into sub-modules since they act as the foundation for a network’s higher level of functional structure. These sub-modules can serve as prospective information sources for a given domain of activity [11].

Life science research now has access to enormous amounts of disease-based histology data with the help of high-throughput sequencing technology. These data are accessible to the public in databases like The Cancer Genome Atlas (TCGA), which has created various high throughput genomic data types for hundreds of samples on different cancer types [12]. To identify the cancer driver genes, it is crucial to specify the genes and the networks connected to them [12]. Finding genes with significant abnormalities and a high frequency of mutations can improve the ability to anticipate how cancer begins and progresses. Finding these cancer-causing driver genes is challenging, and many mutations have not been found using the available databases [13]. Recent studies have shown that, despite some cancer genes having high frequencies of mutation, the majority of cancer genes in most patients have intermediate or low rates of mutation (2–20%) [13]. Therefore, identifying dysregulated pathways and viable targets for therapeutic intervention will require a comprehensive record of mutations in different frequency classes [13]. A recent study shows significant gaps in our knowledge of cancer genes with intermediate frequency. For example, a study of 183 lung adenocarcinomas found that 38% of patients had 3 or fewer of these mutations, and 15% did not have even one mutation that affected any of the ten known hallmarks of cancer [14]. As a result, we cannot fully characterize the expression profiles of all the genes and subsets of genes responsible for the development and spread of cancer. A few pathways, particularly those involved in survival, cell division, differentiation, and genomic preservation, are where cancer genes frequently undergo significant change. So, even for genes with low rates of mutation, it is important to figure out how important they are at the pathway level [15].

The main disadvantage of most of the current methods is that they require extensive filtering of mutation data, which is limited to the most significantly mutated genes and focuses on predefined network modules [16]. These methods may be biased towards identifying gene sets where most of the coverage comes exclusively from highly mutated genes [17, 18]. Although cancer-related genes have been shown to be implicated in multiple pathways, few methods specify the important gene sets where a gene has various mutually exclusive correlations with other genes in various pathways at different mutation frequencies [17–23]. Some methods, such as WITER [19] and driverMAPS [20], are based on the driver genes mutation frequency. These methods are based on the idea that the mutation frequency in driver genes is higher than the background mutation frequency. Some other methods are network-based identification of promoter genes such as HotNet2 [21], NetSig [22], DNmax [23], nCOP [24] and MaxMIF [25]. In these methods, pathways, networks, and mutation frequencies are studied. Some of these network-based methods could identify the number of low-frequency mutated genes.

In this work, in a two-step technique, we proposed two novel algorithms. The first algorithm identifies candidate driver genes in breast cancer with a different frequency of mutations to find the complete mutation pattern. The second algorithm finds the driver modules associated with breast cancer. We used TCGA driver gene information for the first algorithm, which includes a set of low- and high-frequency mutations from thousands of breast cancer patients. Then, we constructed a biological network corresponding to these genes. We defined informative topological and biological features for each node (gene) in the specified network. Our novel machine learning algorithm in this part assigned the score to each node and introduced the high-score genes with expressive connections to breast cancer as candidates for more investigation with the help of the second algorithm. For the second algorithm, we presented a Monte Carlo algorithm to identify the high-score breast cancer-related modules based on the results of the first algorithm. Lastly, we looked at how important our proposed modules were in terms of their role in critical signaling pathways involved in breast cancer. We got a comprehensive insight into breast cancer development when we looked at the breast cancer-related driver modules and driver genes.

## 2 Materials and methods

This part proposed a two-step technique for finding driver genes and modules in breast cancer. In the first step, we presented an unsupervised machine learning algorithm (SPNFSR) for scoring the set of mutated genes in breast cancer. Then, with the help of biological process terms related to genes with high scores, we built the breast cancer network. We identified genes from the breast cancer network with high scores as breast cancer driver genes. In the second step, we presented a Monte Carlo MC-BCM algorithm to find dense clusters with the highest score. Then we identified a set of significant clusters with high p-values based on cancer-related pathways as a set of modules related to breast cancer. The general workflow of the proposed method is illustrated in Figure 1.

**Figure 1:**
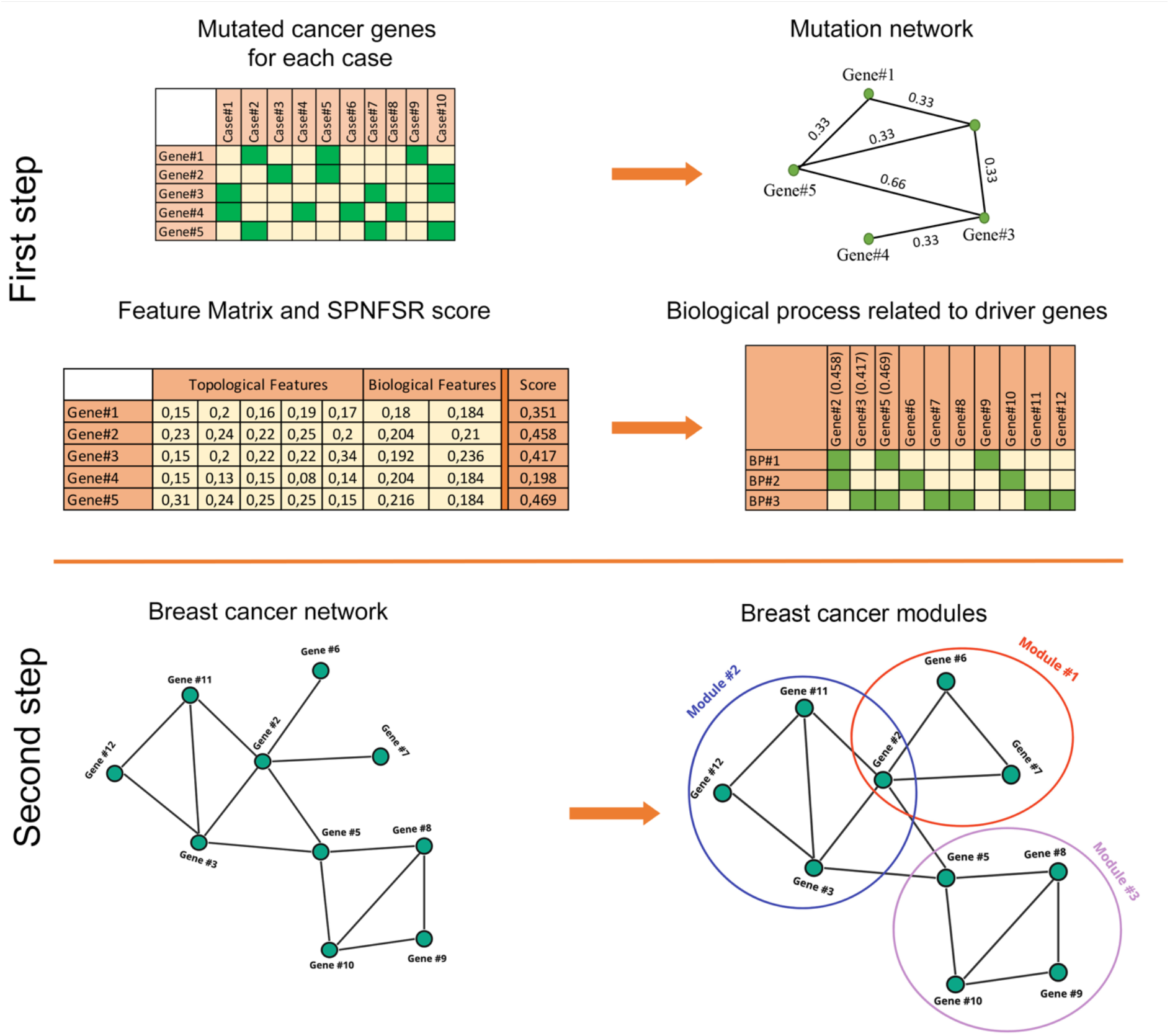
The general workflow of the proposed method. In the first step, we used the information of different patients with breast cancer and their associated mutated genes to create the mutation network. Then, five topological and two biological features are calculated for each node of the mutation network. We used a machine learning method to calculate the suitable scores for each feature. Finally, our algorithm chooses a set of genes with higher scores as a set of mutated genes with valuable information. In the second step, we created the breast cancer network based on the gene relationship to determine the cancer-related modules. We used the information on biological interactions between the high-score genes obtained in the first step to build this network. Then, we used a Monte Carlo algorithm to find breast cancer driver modules in the created weighted network.

### 2.1. Dataset

We used four groups of data for different parts of this study. The first one is the dataset from which we extracted the mutation network. The second one is the dataset from which we extracted biological process annotation related to breast cancer. The third one is the dataset from which we extracted the PPI network. The fourth one is the dataset from which we extracted driver gene data as the benchmark for the evaluation of our result.

#### Mutated Genes

One of the critical steps to identifying driver genes and modules related to breast cancer is to extract significantly mutated genes from breast cancer patients. Researchers have collected numerous datasets on genetic changes in breast cancer patients. One of these datasets with high confidence is TCGA [12]. We downloaded a collection of mutated genes in breast cancer patients and related biological information such as SNV (Single-nucleotide variants) and CNV (Copy number variation) on June 2022. We extracted information on 597 genes with genetic modifications in more than one case.

#### Biological Process

To specify the driver genes and modules related to breast cancer, we used biological process Gene Ontology (GO) terms. Habibi et al [26] identified a list of 62 biological process terms related to cancer. With the help of genes in these GO terms, we expanded the set of mutated genes related to breast cancer. This expansion of genes helps us to have a comprehensive study of other genes that do not have genetic modifications.

#### Protein-Protein Interaction Network

To determine the interaction between genes, we used the PPI network information extracted by Habibi et al [27].

#### Real Driver Genes

Comparing driver gene identification approaches is extremely difficult because there isn’t a comprehensive benchmark for gathering a set of driver genes. In this study, we chose five independent datasets and used them as benchmark sets. The first dataset includes 118 genes, which we presented as “CGCpointMut”. This set contains genes participating in carcinogenesis through point mutations [28]. The second dataset, which is shown as the “Rule” has 124 cancer genes that are included in oncogenes and tumor suppressor genes based on distinctive mutation patterns [15]. The third set contains 288 driver genes with high confidence indicated by at least two frequency-based techniques. This set is denoted by the “HCD” [29]. The fourth set, CTAT, contains 297 driver genes [30]. The fifth set includes 232 driver genes, which are gathered from a mixture of seven driver gene sets. We pointed out this set by the “ShiBench” [31].

### 2.2 Construction of the mutation network

We introduced a mutated network based on 597 mutated breast cancer genes in this work. Suppose that *V* = {*g*_1_,…, *g_n_*} denotes the set of mutated breast cancer genes. Then, assume that 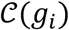 is the set of cases that hold a given mutated gene (*g_i_*). A weighted mutated network *G*= < *E, V, w*> was created by connecting two genes (*g_i_* and *g_j_*) if and only if 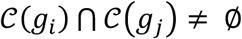. The weight of edge *g_i_g_j_* ∈ *E*, which is indicated by *w*(*g_i_g_j_*), is defined as follows:

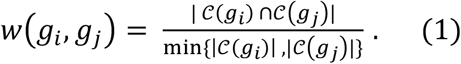

A path between *g_i_* and *g_j_* is defined as a sequence of separate nodes such that an edge of *G* links two consequent nodes. A path’s weight is equal to its edges’ total weights, and a path between two nodes with the least weight is the shortest path from node *g_i_* to node *g_j_* [32]. The weight of the shortest path between two nodes (*g_i_* and *g_j_*) is indicated by *d_w_*(*g_i_, g_j_*).

#### 2.2.1 Topological features for mutation network

We defined the following topological features for each node of the weighed mutation network.

##### Weight

The Weight of node *g_i_* on weighted graph *G* = < *V, E,w* > is defined as follows:

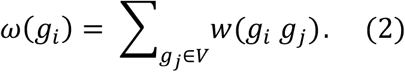

##### Closeness

The Closeness centrality measure for each node, *g_i_*, is defined as follows:

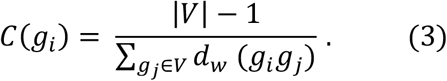

##### PageRank

The score for each node *g_i_* in the network is computed based on all the scores assigned to all nodes *g_j_*, which are joined iteratively as follows:

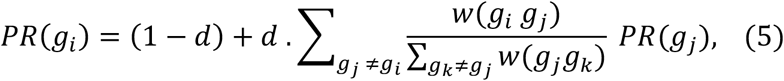

where *d* is a parameter between 0 and 1.

##### Eigenvector

The Eigenvector centrality measure as the amount of influence for a node *g_i_* in the network is defined as follows:

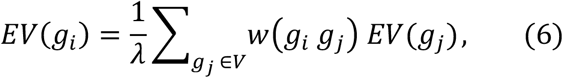

where *λ* is a constant.

##### Entropy

Assume that *ω*(*g_i_*) is the weight of node *g_i_* on weighted network *G* = < *V, E, w* >. The probability distribution vector Π = < *π*_1_,…, *π_|V|_* > is represented on the set of all nodes of the network as follows:

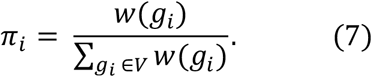

Then, the entropy of weighted network *G* is calculated as follows:

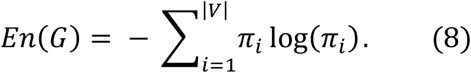

The effect of each node *g_i_* on network entropy is also calculated as follows:

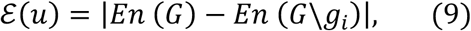

where *G\g_i_* is the weighted network that is created with respect to the removal of node *g_i_* and its linked edges from the network.

#### 2.2.2 Biological features for mutation network

We used the following biological features for each node of the weighed mutation network.

##### SNV

Single nucleotide variants (SNVs) present as germline or somatic point mutations as important drivers of tumorigenesis and cellular proliferation in numerous cancer types.

##### CNV

Copy number variation (CNV) refers to the possibility that the number of copies of a specific DNA segment changes among individuals’ genomes. These structural differences may have come via duplications, deletions, or other modifications and can impact long stretches of DNA.

### 2.3 Machine learning algorithm to identify driver genes associated with breast cancer

In this subsection, we proposed a machine-learning algorithm to compute an effective score for each mutated cancer gene with respect to informative topological and biological features. Then, we extended the high score mutated genes with the help of a set of noteworthy biological processes to detect the candidate driver modules for breast cancer.

#### 2.3.1 SPNFSR algorithm to select top mutated cancer genes

Since there is currently no definitive solution to the challenge of choosing the set of mutated candidate cancer genes, it can be treated as a problem without an exact solution. In order to identify an effective list of mutated cancer genes, we used an efficient unsupervised feature selection algorithm. Assume that *X* = [*x_ij_*]*_m×n_* represents the feature matrix and *x_ij_* represents the j-th feature of the i-th genes as samples. We specified a feature vector 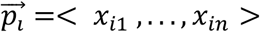 for each sample and determined the column matrix *F_j_* = [*x*_1*j*_,…, *x_mj_*]*^T^* for the j-th feature. We presented the Structure Preserving Nonnegative Feature Self-Representation (SPNFSR) algorithm as an unsupervised machine learning method to compute the score for each mutated gene.

Assume that *S_m×m_* indicates the weighted matrix where *S* = (|*S*| + |*S^T^*|)/2 depicts the similarity of two feature vectors 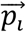 and 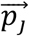. Arranged two identity matrices *R_m×m_*, *Q_n×n_*. Set matrix *L* = (*I* – *S* – *S^T^* + *SS^T^*) and *M* = *X^T^LX*. Assume that *M* = *M^+^* – *M^-^* where *M^+^* = (|*M_ij_*| + *M_ij_*)/2 and *M^-^* = (|*M_ij_*| — *M_ij_*)/2. The components of weighted matrix *W* are iteratively computed as follows:

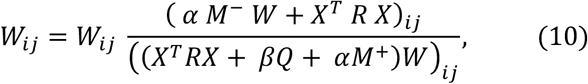

where α, β ≥ 0 and *R* and *Q* as diagonal matrices iteratively updated as follows:

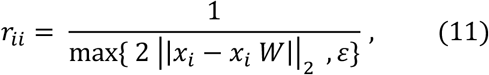

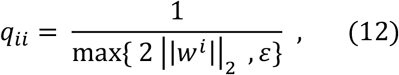

where ε is a small constant. To calculate each feature’s weight, we used 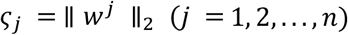 where *w^j^* denotes the j-th row of the weighted matrix *W*. The SPNFSR score was finally determined for each mutated cancer gene *g_i_* as follows:

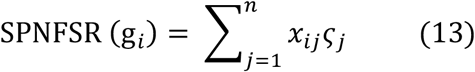

The steps to calculate SPNFSR score for each mutated cancer gene is described in Algorithm 1. In this algorithm, we considered that *α* = 0.1, *β*=0.1, *γ* = 5 and *t* = 10 respectively.

##### Algorithm 1 Structure Preserving Nonnegative Feature Self-Representation (SPNFSR)

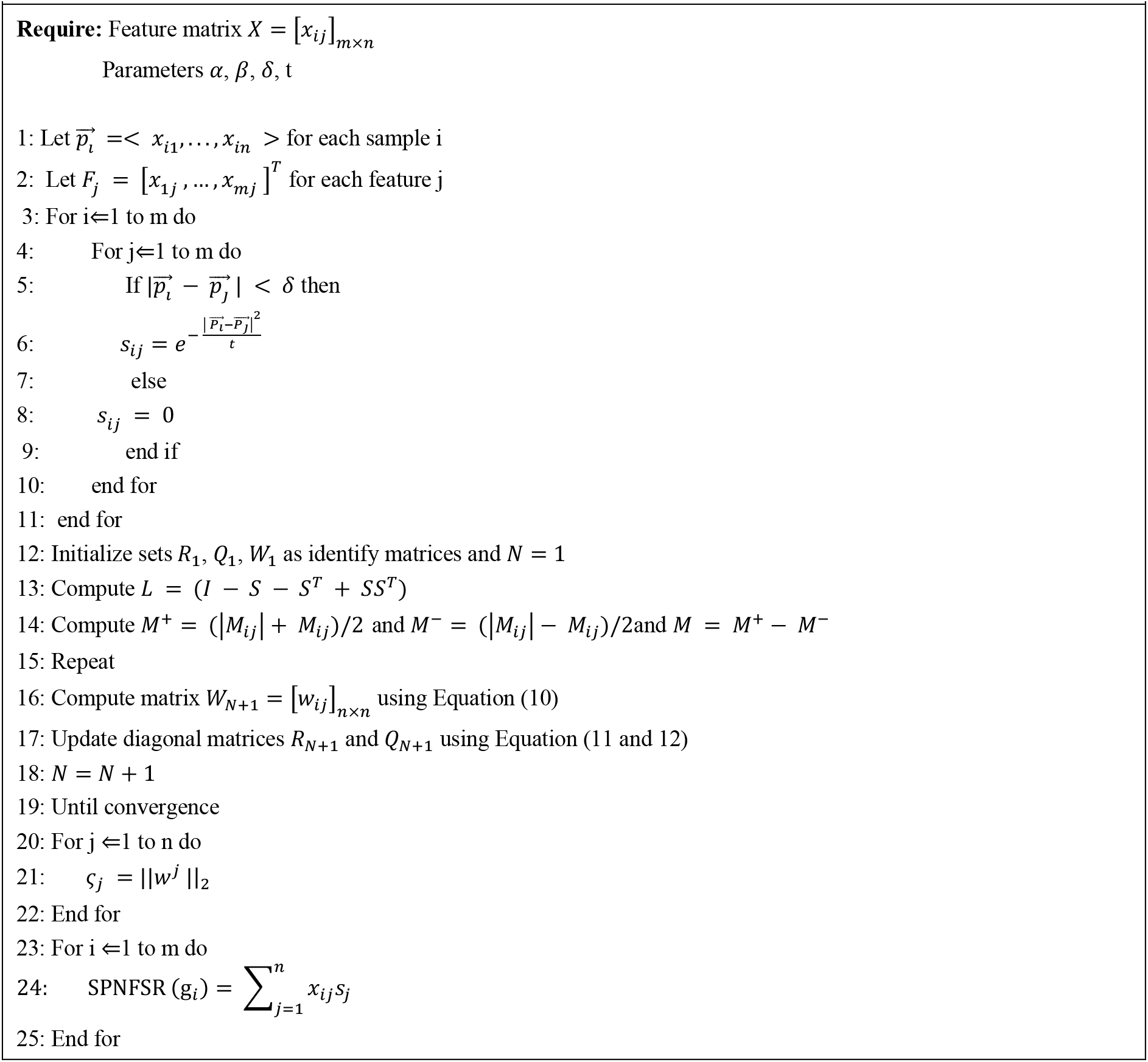

#### 2.3.2 Construction of the breast cancer network

Suppose that *M_G_* = {*g*_1_,…, *g_m_*} is a set of *m* mutated genes under study, where each gene’s *SPNFRS*(*g_i_*) score has been calculated in the previous subsection. We selected a set of genes with a high score as the top genes related to breast cancer, and we denoted this set with the *T_G_*. In set *T_G_*, the total score of genes includes 25% of the total scores. Let *BP* = {*P*_1_…, *P_k_*} is a set of biological processes related to breast cancer, which includes at least one of the *T_G_* genes. We defined the set of breast cancer network genes as follows:

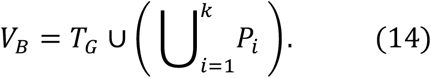

Then, we described the weight of each gene in this network as follows:

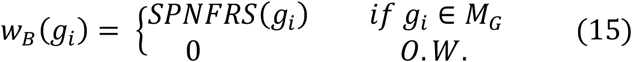

We introduced the network *G_B_* = < *V_B_, E_B_, w_B_* > as a weighted breast cancer network, where each edge *g_i_g_j_* ∈ *E_B_* represents an interaction between genes *g_i_* and *g_j_*. Then, we defined a set of genes in *V_B_* with a higher weight than zero as the set of driver genes related to breast cancer. In other words, we introduced the shared genes between *V_B_* and *M_G_* as the breast cancer driver genes network.

### 2.4 MC-BCM Algorithm to Identify Breast Cancer driver modules

Suppose *G_B_* = < *V_B_, E_B_, w_B_* > is a breast cancer network. The density of each cluster *H* = < *V_B_*(*H*), *E_B_*(*H*), *w_B_*(*H*) > as a subnetwork of the graph *G_B_* is defined as follows:

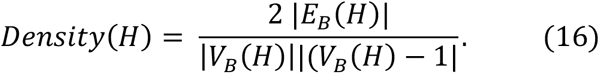

The density value of each cluster is between 0 and 1. The dense subgraphs have a closer density value of 1. The average weight of each cluster’s nodes is also called that cluster’s weight.

Two nodes of the graph are called neighbors if there is an edge between those two nodes. We denoted the set of all neighbors of the node *v* with the symbol *N*(*v*) and called it the neighborhood of *v*. Let *H* be a cluster of the network, then, the neighborhood of cluster *H* is defined as follows:

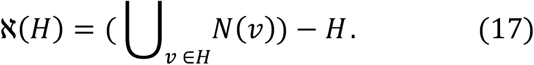

We called cluster *H*_1_ as a neighbor of cluster *H* if the number of nodes of both clusters is equal and *V*(*H*_1_) – *V*(*H*) = {*v*} so that 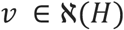. We denoted the set of all clusters that are neighbors to cluster *H* with *Neig*(*H*). Suppose that *P* = {*p*_1_,…, *p_M_*} is a collection of significant pathways related to breast cancer. Also, suppose that *H* is a candidate cluster for breast cancer. We used the Gene Set Enrichment Analysis (GSEA) formula to calculate the p-value of cluster *H* with respect to the pathway *P_i_* as follows:

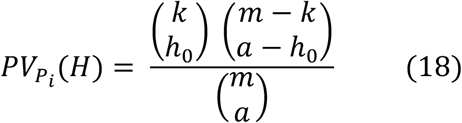

where *m* is the number of genes in the breast cancer network, *a* is the number of genes in the cluster, *k* is the number of genes in the pathways *p_i_*, and *h*_0_ is the number of cluster genes in the assumed pathway.

A cluster was defined as a breast cancer driver module if it had a significant p-value, a high density, and a high mean score. We presented the MC-BCM (Monte Calo algorithm for breast cancer driver module) to find driver modules related to breast cancer. This algorithm has two steps. In the first step, with the help of the Monte Carlo algorithm, we tried to find the dense subgraphs of the breast cancer network. In the second step, we proposed two filters for the set of discovered clusters. The first filter is based on the weight of dense clusters, and the second filter is based on the p-value of the cluster corresponding to a set of significant pathways related to breast cancer. In Algorithm 2, we showed the details of finding high-score dense driver modules.

#### Algorithm 2 Monte Carlo algorithm for breast cancer driver module (MC-BCM)

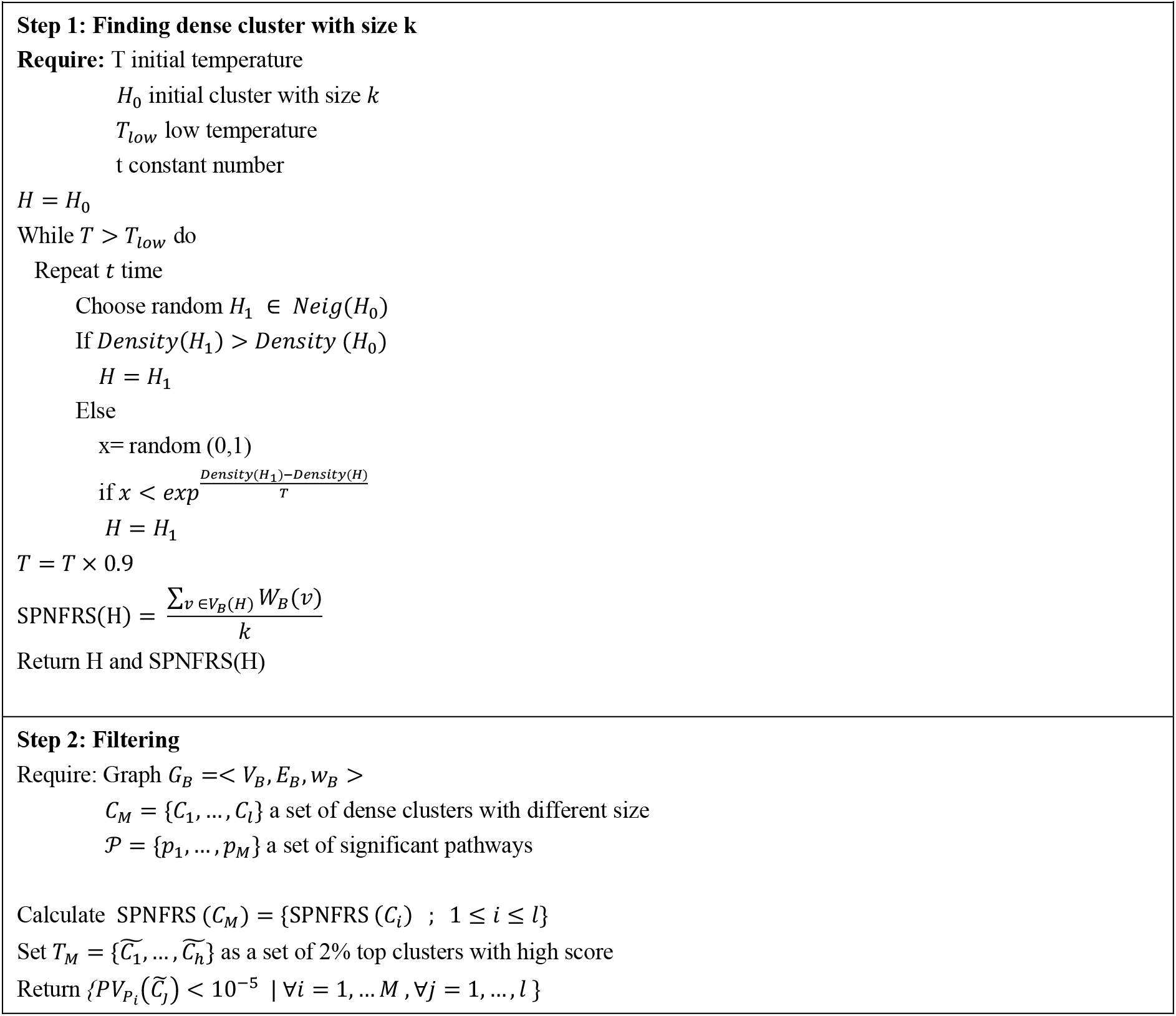

Suppose that *C_M_* = {*C*_1_,…, *C_l_*} is a set of clusters with high-density and high scores obtained by performing the MC-BCM with different *k* (4 ≤ *k* ≤ 10) and for different initial clusters. We selected 2% of the clusters with high scores as the set of candidate clusters. Then, with the help of the GSEA formula (18), we computed the p-value of each candidate cluster for each significant pathway. Now, the set of clusters with a p-value less than 10^-5^ is introduced as the breast cancer modules.

## 3. Results

The results of this study have two parts. The first part shows the results for identifying mutated driver genes related to breast cancer. The second part shows the results for determining functional modules related to breast cancer. Each of these parts has been evaluated and studied separately.

### 3.1 Results of SPNFSR algorithm in finding driver genes related to breast cancer

From 12,987 cases with genomic changes reported on TCGA in June 2022, 944 cases were involved in breast cancer. This work has studied the changes in 597 genes in these cases. Among these genes, TP53 has the highest genomic changes in more than 22% (209/944) of the studied breast cancer cases. After that, MUC16, MAP3K1, CDH1, and KMT2C genes, with more than 17% (167/944), 12% (116/944), 12% (114/944), and 10% (97/944), respectively, have the most significant number of breast cancer cases with genomic alterations. Notably, more than 67% (401/597) of the studied mutated genes were observed in fewer than 10 cases. The study of some of these genes with low frequency showed that the genomic changes in these genes are essential in the cancer process. Therefore, identifying these genes is critical in determining breast cancer’s pathology. In other words, studying the number of cases with high-frequency genomic changes is not enough to study breast cancer. Thus, in this study, we are trying to find a set of important mutated genes related to breast cancer, which, in addition to the frequency of mutations, also pay attention to other biological characteristics. In this work, we have considered a two-step approach to identify driver genes related to breast cancer. We tried to score genes based on topological and biological features in the first step. Since finding such a score for the set of genes with genetic alterations related to breast cancer is still an open problem, we solved this problem with the SPNFSR as an unsupervised machine-learning method. In this step, we extracted a set of top genes with the highest score. In the second step, we prepared a list of genes with the help of biological processes related to breast cancer in which this set of top genes participated. Then, we select a set with 262 mutated genes with the highest score as a set of driver genes related to breast cancer.

#### 3.1.1 Evaluation of SPNFSR algorithm

We studied different topological and biological features in this study. With the help of the SPNFSR algorithm as a feature selection algorithm, we have selected a list of significant topological and biological features. We introduced five topological and two biological features. These topological and biological features contain meaningful information about the network. Table 1 shows the weight of each of these features.

**Table 1.** The score is calculated by the SPNFSR algorithm for topological and biological features.

| Features | Score |
| --- | --- |
| Weight | 0,136 |
| Closeness | 0,074 |
| PageRank | 0,1 |
| Eigenvector | 0,137 |
| Entropy | 0,08 |
| SNV | 0,074 |
| CNV | 0,059 |

To evaluate the SPNFSR algorithm, we created a simulated dataset using genes related to prostate cancer. For this purpose, we extracted the set of genes related to the growth factor pathway from the mutated genes related to prostate cancer, which were introduced in the TCGA. This gene set contains genes that increase cell proliferation and oncogenes, such as AR. Then, we added 100 genes to the previous set, including 20 genes related to and 80 genes unrelated to prostate cancer. From the collection of 200 defined cases, we assigned a case for each of these genes related to prostate cancer at a rate of 40% and for unrelated genes at a rate of 20%. We also selected a number between 15 and 20 SNVs for each cancer-related gene and between 10 and 20 for each unrelated gene. Similarly, for each cancer-related gene, the CNV values for each case are between 5 and 10, and the CNV values for each non-cancer-related gene case are between 0 and 10. Then, we used the SPNFSR algorithm and assigned a score to each simulated data. We obtained 23 genes with the highest score as the algorithm’s output. Among the 20 genes related to prostate cancer, 15 were selected as genes with a high score.

To evaluate the output of the SPNFSR algorithm, we used the GSEA formula (18) to obtain p-values for the prostate cancer gene set. In this formula, *m* indicates the number of simulated genes (*m* = 100), *a* is the number of genes that we have chosen as the top gene (*a* = 23), and *k* is the number of genes related to prostate cancer (*k* = 20). The number of common genes between our selected top genes and genes related to prostate cancer is equal to *h*_0_ = 15. The p-value for this data set driven by our selected gene set equals 1.8 * *E^-8^*. This p-value shows that our set of top genes was not chosen randomly or by chance.

#### 3.1.2 Evaluation of high-score selected genes based on SPNFSR

Finding a new set of genes with important properties, even if they have moderately or infrequently mutated, guides us to new informative modules. To evaluate the SPNFSR algorithm’s performance, we apply SPNFSR to a set of mutations collected from TCGA. The proposed machine learning method calculated the score for each candidate mutated gene in this set. The total score of 597 candidates’ mutated genes is 734.24, and the average score of these candidate genes is 1.23. From 597 candidate genes, PIK3CA has the highest score of 3.18, and DDIT3 has the lowest score of 0.49. We have selected a list of 100 informative genes with high scores that include 25% of the total scores (734.24 × 25%) as the gene set with the highest score. Figure 2 shows the heat map of these genes that are arranged from large to small based on the SPNFSR score. The value of each feature and the frequency of each case are also shown in this heat map. In this figure, the frequency of the cases shows that our candidate high-score gene set includes genes with high frequency, such as TP53 and PIK3CA, and also genes with low frequency, such as HOXA9 and DDB2.

**Figure 2:**
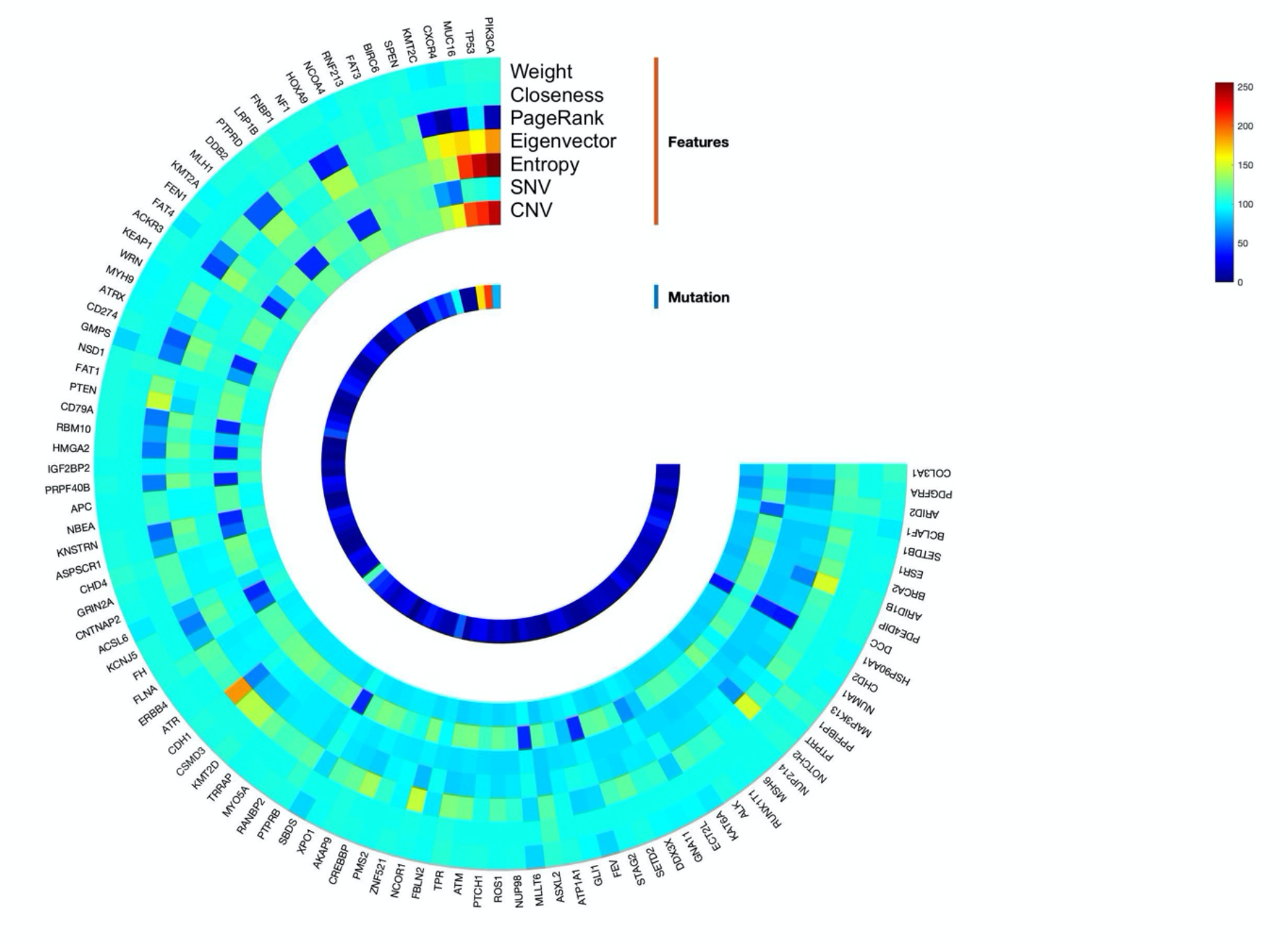
heat map of high score genes that are arranged from large to small based on the SPNFSR score.

In this study, we considered genes with fewer than 10 cases as low-mutated genes. Among the 100 high-score selected genes, 30 genes have low-frequency mutations. Table 2 shows these genes and their corresponding scores. In the following, we described a correlation between some of these genes with the highest score among these 30 genes and breast cancer to show their importance even with the low frequency of mutation.

**Table 2.**
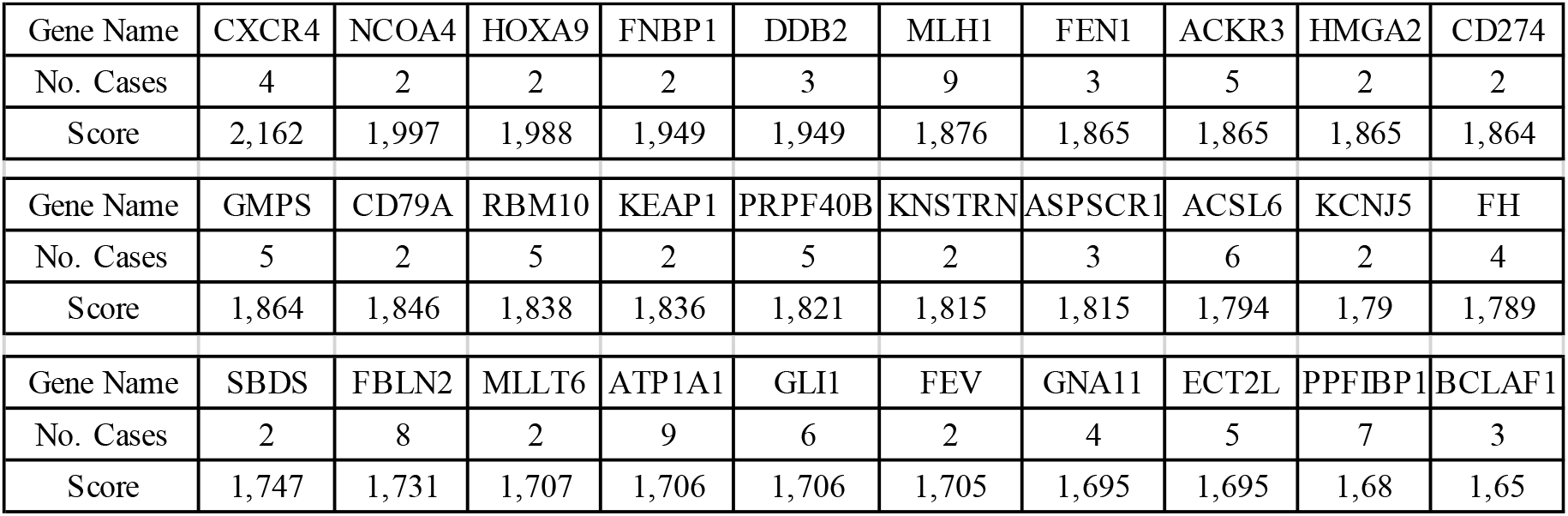
A list of 30 low-frequency mutated genes with high-score.

##### CXCR4

Since identifying CXCR4 in metastatic breast cancer, many studies have established CXCR4 as a critical regulator of primary and metastatic breast cancer [33]. It seems that the CXCR4 signaling pathway facilitates breast cancer through a broad range of mechanisms, such as proliferation, survival, and angiogenesis at primary and metastatic sites [33].

##### NCOA4

Recent studies showed that two isoforms of NCOA4, NCOA4α, and NCOA4β, have been functionally evaluated in breast cancer [34]. NCOA4α inhibits proliferation, whereas NCOA4β increases proliferation in breast cancer cells [34].

##### HOXA9 and HMGA2

The TET family starts the DNA’s demethylation process and is linked to carcinogenesis in several malignancies. Many studies have identified upstream and downstream activators and effectors of TET1 in breast cancer [35]. They showed that HMGA2 generates TET1. TET1 binds and demethylates its own and HOXA promoters by improving its expression and stimulating HOXA gene expressions such as HOXA7 and HOXA9. Both TET1 and HOXA9 suppress growth and metastasis in breast tumors from mouse xenografts. The genes containing the HMGA2–TET1–HOXA9 pathway are regulated in breast cancer and include a predictive signature for patient survival [35]. These results showed the effect of the HMGA2–TET1–HOX signaling pathway in the regulation of human breast cancer [35].

##### FNBP1

The FNBP1 has a critical role in tumorigenesis and progression and is significantly expressed in different tumor tissues. A high expression level of FNBP1 is correlated with a better prognosis in breast cancer cells. It correlates with diverse immune infiltrating cells and various immune gene markers in invasive breast carcinoma. Authors in [36] revealed that FNBP1 might significantly influence breast cancer prognosis and immune infiltration levels as a key biomarker. They showed that the expression level of FNBP1 contributes to assessing metastatic and non-metastatic tumors and the immune escape mechanism because of its elevated expression level at immune checkpoints [36].

##### DDB2

Recent studies showed DDB2 is involved in DNA repair and cell cycle regulation. Functional analysis of DDB2 expression demonstrated that DDB2 negatively regulates the expression of the SOD2 gene in breast cancer cells. DDB2 activity occurs at different breast cancer stages, such as proliferation, survival, and migration [37].

##### MLH1

Mismatch repair deficiency contributes to breast cancer progression into progressive stages and assists cancer cells to survive against chemotherapy drugs. Recent analysis indicated that MLH1 is essential in preserving the DNA repair system, and loss of this mismatch repair protein could cause negative results in breast cancer [38].

##### FEN1

The FEN1 is recognized as a key factor in DNA replication and telomere maintenance. Elevated levels of FEN1 expression have been reported to be associated with cancer progression and promote breast cancer cell proliferation [39].

#### 3.1.3 Evaluation of driver genes related to breast cancer

In this study, we collected 61 biological processes GO terms that are associated with breast cancer. All of these 61 biological processes have at least one gene from the set of 597 understudied genes. 1193 genes have participated in this set of biological processes. Among 1193 genes, we have selected 262 genes with the highest score as driver genes related to breast cancer. One of the significant challenges of driver gene identification algorithms is that there is no specific and accurate benchmark to compare these methods, which has made evaluating driver gene identification algorithms extremely difficult. In this work, to assess the set of driver genes identified by the SPNFSR algorithm, we calculated the p-values for each benchmark set, CGCpointMut, Rule, HCD, CTAT, and ShiBench, respectively. Let *a* be the number of driver genes recognized by our algorithm (*a* = 262), *m* be the number of all the understudied genes (597 genes), and *k* be the cardinality of the union of driver genes between the benchmark set and understudied genes, and *h*_0_ be the cardinality of intersection between driver genes identified by our algorithm and the benchmark set. The probability of being placed in the set of shared genes with a size *h*_0_ is calculated from (18). Table 3 displays the p-values for each of the benchmark sets, with values of *k* and *h*_0_. According to the p-values in Table 3, our set was not chosen at random. For better evaluation of the driver genes that are identified by our algorithm and the driver genes identified by other algorithms for breast cancer, we need to define statistical measures to determine the agreement between them and evaluate them with benchmark sets. We used the following known criteria:

**Table 3.** The p-values for candidate driver genes in benchmark sets. The number of driver genes in the benchmark set among the 597 mutated understudied genes is represented by *k*, and the number of driver genes recognized by our algorithm in each benchmark set is represented by *h*_0_.

|  | CGCpointMut | CTAT | HCD | Rule | ShiBench | Union |
| --- | --- | --- | --- | --- | --- | --- |
| k | 101 | 166 | 149 | 104 | 191 | 483 |
| $h_0$ | 55 | 97 | 88 | 64 | 108 | 228 |
| p-value | 0,0052 | 3,17E-06 | 6,13E-06 | 2,77E-05 | 6,21E-06 | 1,42E-04 |

True Positive (*TP*) indicates the genes correctly identified as the driver genes. The True Negative (*TN*) shows the genes correctly identified as the non-driver genes. The False Positive (*FP*) represents the genes incorrectly identified as the driver gene, and the False Negative (*FN*) indicates the genes incorrectly identified as the non-driver gene. The evaluation parameters of Precision (*Pre*), Recall (*Rec*), and F-measure (*F*) are also defined as follows respectively.

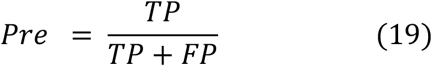

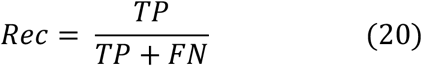

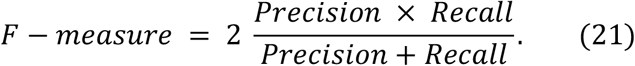

Recently, different methods with different approaches have been proposed to identify cancer driver genes. For more investigation, we compared the driver genes identified by DriverML, Dmax HotNet2, MaxMIF, nCOP, NetSig, and WITER algorithms and our algorithm. Table 4 shows that our algorithm has an acceptable agreement with other methods, especially DriverML, MaxMIF, Dmax, and WITER. Since the WITER and DriverML methods are statistical frequency-based methods and the Dmax and MaxMIF are network-based methods, it can be concluded that our algorithm has excellent agreement with different approaches and methods. To further study the difference between our algorithm and these algorithms, we focused on driver genes that are recognized by our algorithm, but other algorithms did not confirm them. We found that among the 262 driver genes recognized by our algorithm, there are eight genes (KMT2C, KMT2A, ACKR3, CD79A, KMT2D, ASXL2, NSD3, PIM1) that were confirmed by at most two algorithms. In the following, we study the association and role of these eight genes in breast cancer.

**Table 4.** Comparison of the performance of our algorithm with other algorithms with different approaches. The high values are shown in bold.

|  | TP | TN | FP | FN | Pre | Rec | F |
| --- | --- | --- | --- | --- | --- | --- | --- |
| Dmax-SPNFRS | 253 | 17 | 320 | 9 | 0,442 | **0,97** | **0,606** |
| DriverML-SPNFRS | 248 | 21 | 316 | 14 | 0,44 | 0,95 | 0,6 |
| HotNet2-SPNFRS | 22 | 307 | 30 | 240 | 0,423 | 0,08 | 0,14 |
| MaxMIF-SPNFRS | 249 | 19 | 318 | 13 | 0,439 | 0,95 | 0,601 |
| nCOP-SPNFRS | 83 | 289 | 48 | 179 | 0,634 | 0,32 | 0,422 |
| NetSig-SPNFRS | 198 | 46 | 291 | 64 | 0,405 | 0,76 | 0,527 |
| WITER-SPNFRS | 245 | 31 | 306 | 17 | **0,445** | 0,94 | 0,603 |

##### PIM1

Due to the poor prognosis of triple-negative breast tumors and the scarcity of identified potential targets, chemotherapy is still the backbone of treatment. In triple-negative breast cancers, MYC was discovered to be overexpressed compared to other subtypes, particularly in those resistant to chemotherapy, but suppression has proven difficult to achieve [40]. Recently, the cooperation of PIM1 and MYC was identified as involved in the migration, cell proliferation, and apoptosis of triple-negative breast cancers. Inhibition of PIM1 can facilitate the apoptosis of tumor cells and improve sensitivity to chemotherapy. Therefore, PIM1 could be a promising target in triple-negative breast cancers [40].

##### NSD3

Researchers in [41] proposed that NSD3 is frequently dysregulated in human cancers. They reported that NSD3-induced methylation of H3K36 is critical for breast tumor initiation and metastasis. In patients with breast cancer, overexpression of NSD3 was associated with recurrence, metastasis, and a low chance of survival. This study showed that mice carrying primary and metastatic breast tumors with elevated NSD3 expression showed sensitivity to NOTCH inhibition. This research revealed the critical role of NSD3 in the modulation of NOTCH-dependent breast tumor progression and recommended targeting the NSD3–NOTCH signaling in aggressive breast cancer [41].

##### ASXL2

Researchers in [42] proposed that Estrogen receptor alpha (ERα) has a crucial role in breast carcinogenesis by associating with various cellular factors. They investigated the involvement of ASXL2 in ER activation and breast carcinogenesis because of its selective expression in ER-positive breast cancer cells. They showed that the ability of cells to proliferate and the size of xenograft mice’s tumors were affected by ASXL2 knockdown [42]. This research revealed that ER-positive individuals had increased ASXL2 expression and suggested that ASXL2 might be a new prognostic indicator for breast cancer.

##### KMT2

Recent studies on KMT2 mutations in humans indicate that these genes are frequently mutated in various cancers, such as breast cancer. Most cancers with KMT2 mutations harbor at least one wild-type allele; it is also important to characterize the function of wild-type KMT2 in malignant versus pre-malignant cells [43]. Many studies on KMT2A and KMT2D have shown that wild-type proteins could be needed for cancer cells to stay alive. Therefore, they could be used as therapeutic targets. Researchers in [44] proposed that breast cancers with estrogen receptor (ER) positive status typically contain activating mutations in the PIK3CA gene, which codes for PI3K. The therapeutic efficacy of PI3K inhibitors, which are now in late-stage clinical development, is constrained by a significant compensatory increase in ER-dependent transcription. They discovered that in breast cancer clinical data, PI3K inhibition promotes an open chromatin state at the ER target loci. KMT2D is necessary for the recruitment and activation of ER, FOXA1, and PBX1. While PI3K inhibition boosts KMT2D activity, AKT binds to and phosphorylates KMT2D, reducing methyltransferase activity and ER function. This research revealed that hybrid therapies consisting of PI3K inhibitors and KMT2D inhibitors may be more effective than using PI3K inhibitors alone. Researchers in [45] proposed that KMT2C is by far the most typically mutated gene in TCGA breast cancer samples. Most of the KMT2C mutations are truncation or missense mutations. The mutations appear in all breast cancer subtypes with a fairly even distribution. They suggested that KMT2C could be a breast cancer tumor suppressor and a candidate regulator of H3K4me in breast cancer tumors. Recent studies show that mutations in the KMT2A gene are prevalent in an extensive range of solid tumors, including colon, lung, bladder, and breast cancers. Most KMT2A mutations are frameshift mutations that lead to the early termination of translation. Therefore, most cancers, like breast cancer, still have at least one wild-type KMT2A allele [45].

##### ACKR3

Researchers in [46] reported the importance and role of ACKR3 in different cancers. This chemokine receptor is overexpressed in multiple cancer types, such as breast cancer. It has been involved in tumor cell proliferation and migration, tumor angiogenesis, or resistance to drugs, thus having a significant effect on cancer progression and metastasis [45]. They focused on the clinical significance and probable mechanisms underlying the pathologic role of ACKR3 in breast cancer. They discussed its possible relevance and possible therapeutic target and predictive factor.

##### CD79A

Signaling pathways perform an essential role in the analysis of cancer pathogenesis. According to the latest studies, CD19, CD22, and CD79A were focused on the B cell receptor signaling pathway [47]. These genes encode a cell surface molecule that creates the antigen receptor of B-lymphocytes to reduce the threshold for antigen receptor-dependent stimulation. These differentially expressed genes were closely associated with potential therapeutic targets for triple-negative breast cancer [47].

We also compared the performances of DriverML, Dmax, HotNet2, nCOP, NetSig, WITER, and SPNFRS algorithms in identifying the reported driver genes. The set of genes reported in CGCpointMu, CTAT, HCD, Rule, and ShiBench has been used as the driver gene benchmark. Table 5 shows the values of all the statistical scores by comparing the driver genes identified by these algorithms with the benchmark sets.

**Table 5.** The values of all the statistical scores for different algorithms on different benchmarks. The high values are shown in bold.

|  | TP | TN | FP | FN | Pre | Rec | F |
| --- | --- | --- | --- | --- | --- | --- | --- |
| Dmax-CGCpointMut | 98 | 23 | 475 | 3 | 0,171 | **0,97** | 0,29 |
| DriverML-CGCpointMut | 96 | 30 | 468 | 5 | 0,17 | 0,95 | 0,288 |
| HotNet2-CGCpointMut | 13 | 459 | 39 | 88 | 0,25 | 0,128 | 0,169 |
| MaxMIF-CGCpointMut | 96 | 27 | 471 | 5 | 0,169 | 0,95 | 0,287 |
| nCOP-CGCpointMut | 30 | 397 | 101 | 71 | 0,229 | 0,297 | 0,258 |
| NetSig-CGCpointMut | 82 | 91 | 407 | 19 | 0,167 | 0,811 | 0,277 |
| WITER-CGCpointMut | 92 | 39 | 459 | 9 | 0,166 | 0,91 | 0,282 |
| SPNFRS-CGCpointMut | 55 | 291 | 207 | 46 | **0,209** | 0,544 | **0,303** |
| Dmax -CTAT | 161 | 21 | 412 | 5 | 0,28098 | **0,9699** | 0,436 |
| DriverML-CTAT | 158 | 27 | 406 | 8 | 0,28014 | 0,9518 | 0,433 |
| HotNet2-CTAT | 11 | 392 | 41 | 155 | 0,21154 | 0,0663 | 0,101 |
| MaxMIF-CTAT | 158 | 24 | 409 | 8 | 0,27866 | 0,9518 | 0,431 |
| nCOP-CTAT | 49 | 351 | 82 | 117 | 0,37405 | 0,2952 | 0,33 |
| NetSig-CTAT | 129 | 73 | 360 | 37 | 0,2638 | 0,7771 | 0,394 |
| WITER-CTAT | 152 | 34 | 399 | 14 | 0,27586 | 0,9157 | 0,424 |
| SPNFRS-CTAT | 97 | 268 | 165 | 69 | **0,37023** | 0,5843 | **0,453** |
| Dmax -HCD | 142 | 19 | 431 | 7 | 0,24782 | **0,953** | 0,393 |
| DriverML-HCD | 139 | 25 | 425 | 10 | 0,24645 | 0,9329 | 0,39 |
| HotNet2-HCD | 7 | 405 | 45 | 142 | 0,13462 | 0,047 | 0,07 |
| MaxMIF-HCD | 139 | 22 | 428 | 10 | 0,24515 | 0,9329 | 0,388 |
| nCOP-HCD | 46 | 365 | 85 | 103 | 0,35115 | 0,3087 | 0,329 |
| NetSig-HCD | 116 | 77 | 373 | 33 | 0,23722 | 0,7785 | 0,364 |
| WITER-HCD | 134 | 33 | 417 | 15 | 0,24319 | 0,8993 | 0,383 |
| SPNFRS-HCD | 88 | 276 | 174 | 61 | **0,33588** | 0,5906 | **0,428** |
| Dmax -Rule | 101 | 23 | 472 | 3 | 0,17627 | **0,9712** | 0,298 |
| DriverML-Rule | 100 | 31 | 464 | 4 | 0,1773 | 0,9615 | 0,299 |
| HotNet2-Rule | 9 | 452 | 43 | 95 | 0,17308 | 0,0865 | 0,115 |
| MaxMIF-Rule | 100 | 28 | 467 | 4 | 0,17637 | 0,9615 | 0,298 |
| nCOP-Rule | 31 | 395 | 100 | 73 | 0,23664 | 0,2981 | 0,264 |
| NetSig-Rule | 83 | 89 | 406 | 21 | 0,16973 | 0,7981 | 0,28 |
| WITER-Rule | 97 | 41 | 454 | 7 | 0,17604 | 0,9327 | 0,296 |
| SPNFRS-Rule | 63 | 296 | 199 | 41 | **0,24046** | 0,6058 | **0,344** |
| Dmax -ShiBench | 178 | 21 | 395 | 5 | 0,31 | **0,9727** | 0,471 |
| DriverML-ShiBench | 174 | 26 | 390 | 9 | 0,308 | 0,9508 | 0,466 |
| HotNet2-ShiBench | 14 | 370 | 38 | 177 | 0,269 | 0,0733 | 0,115 |
| MaxMIF-ShiBench | 176 | 24 | 391 | 8 | 0,31 | 0,9565 | 0,468 |
| nCOP-ShiBench | 56 | 333 | 75 | 135 | **0,427** | 0,2932 | 0,348 |
| NetSig-ShiBench | 152 | 71 | 337 | 39 | 0,31 | 0,7958 | 0,447 |
| WITER-ShiBench | 174 | 34 | 377 | 14 | 0,315 | 0,925 | 0,47 |
| SPNFRS-ShiBench | 107 | 253 | 155 | 84 | 0,408 | 0,5602 | **0,472** |

Table 5 shows that the Dmax algorithm has identified 573 genes among the 597 genes as breast cancer driver genes, so the precision value in all the studied benchmark sets is close to 100. However, many of these mutated genes are not necessarily breast cancer driver genes. In other words, the FP value of this algorithm is high, and this algorithm does not show an informative analytical representation.

The DriverML, MaxMIF, WITER, and NetSig algorithms have identified 564, 567, 551, and 489 genes among 597 mutated genes as driver genes related to breast cancer, respectively, and many of these genes are *FP*. In other words, the *Re* value of these algorithms does not show an acceptable display (based on F-measure) compared to the SPNFRS algorithm. Among the studied algorithms, the HotNet2 algorithm identified only 52 genes among the 573 mutated genes, but many of these genes were identified incorrectly. Therefore, compared to the low number of genes predicted by the HotNet2 algorithm, it has a high *FP* value in most benchmark sets. Therefore, this algorithm’s amount of *Pre* and *Re* is low.

The nCOP algorithm identified 131 driver genes out of 597 genes and showed an acceptable performance compared to other algorithms. Among these 131 genes, a high number of them have been correctly identified in each benchmark set (*TP*), and as a result, it shows a better representation in *Pre*. However, the SPNFRS algorithm has significant superiority compared to nCOP and other algorithms in *Pre*. The fact that the SPNFRS algorithm has a higher *Pre* is related to the strength of our approach in F-measure across all benchmark sets.

### 3.2. Evaluation of MC-BCM algorithm and candidate driver modules

One of the essential goals of this study is to find modules related to breast cancer. In this study, we introduced dense clusters with a high mean score as a candidate driver module related to breast cancer. For this purpose, we have considered the genes in 61 biological processes GO terms related to cancer, which include 100 genes with high scores. This set of biological processes contains 1193 genes. Among these 1193 genes linked to breast cancer, 262 genes were introduced and evaluated as breast cancer driver genes in the previous section. The breast cancer network is considered a weighted network whose nodes contain 1193 genes and whose edges correspond to their interactions [48]. Among the 1193 nodes of the breast cancer network, 262 nodes are weighted, corresponding to the score obtained by the SPNFRS algorithm, and we considered the weight of the other nodes to be zero. To identify the candidate set of driver modules related to breast cancer, we used the MC-BCM algorithm, then filtered the clusters based on the average scores of each cluster. For this purpose, we ran the Monte Carlo algorithm to find dense clusters with sizes of 6, 7, 8, 9, and 10, respectively and identified 5000 dense clusters for each size. We calculated the average score of each cluster’s nodes as that cluster’s score. Among the 25,000 obtained dense clusters, we extracted 2% of the clusters with the highest average score. This set contains 500 dense clusters with a high score, which we considered as the candidate driver module set. In this section, we evaluated the algorithm’s performance in finding dense clusters with a high score. Then we extracted a collection that includes 75 modules based on the p-value of the pathways participating in that module.

#### 3.2.1 Evaluation of MC-BCM algorithm based on random sets

To evaluate the performance of the MC-BCM, we compared the set of clusters extracted by the MC-BCM algorithm with the random set of clusters. Suppose that *C* = {*c*_1_, *c*_2_,…., *c_n_*} is a set of clusters obtained by the MC-Mod BCM. The number of clusters from the set *C* whose density is greater than the threshold *t* is represented by *N_D_*, and the number of clusters from the set *C* whose nodes’ average score is greater than the threshold *t*_0_ is denoted by *N_s_*. We randomly extracted 1000 sets from the cluster *C^j^* where *j* = (1,…, 1000). Each set of clusters *C^j^* contains *n* clusters with the same size as the clusters of set *C*. With *N_Dj_*, we displayed the number of clusters from the set *C^j^* that had a higher density than the threshold *t*. With *N_sj_*, we demonstrated the number of clusters from the set *C^j^* with an average score of nodes greater than the threshold *t*_0_. Assume 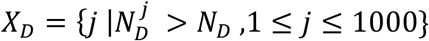 and 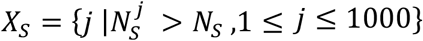. Here, *X_D_* shows the number of sets of random clusters that have better performance than our algorithm based on density, and *X_s_* shows the number of sets of random clusters that have better performance than our algorithm based on the average score of nodes. The null hypothesis *H*_0_ is that our cluster set is insignificant. The alternative hypothesis *H*_1_ is that our set of clusters is significant. We use Exceeding Value (*EV*) to evaluate our selected set of clusters [48].

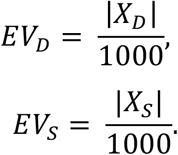

If the *EV* is less than a defined threshold, we reject *H*_0_, and if the *EV* is equal to zero, the results are highly significant. The results of our investigations for different thresholds *t* and *t*_0_ show that the values of *EV_D_* and *EV_S_* equal zero. Therefore, the results of our algorithm have better performance both in terms of density and scores for all sets of random clusters.

#### 3.2.2 Evaluation of modules related to breast cancer

Many studies have tried to identify signaling pathways involved in breast cancer, and due to the disruption of these signaling pathways, cancer development is a complex process. To determine the significant modules related to breast cancer, we have collected a set of signaling pathways involved in breast cancer [50]. These signaling pathways affect cell responses such as survival, proliferation, migration, differentiation, and apoptosis and promote breast cancer [50]. In the following, we summarize these signaling pathways and their relationship with breast cancer.

##### 1. Ras/Raf signaling pathway

Targeting the Ras/Raf signaling pathway and its upstream activators in breast cancer has received much interest. Recent studies indicate that this pathway is abnormally activated in breast cancer with high frequency, particularly by upstream activation of EGFR [50]. Abnormal activation of Ras proteins is the primary inspiration of oncogenes that have an essential role in the main signaling pathway in cancer, and mutation of Ras proteins causes cancer development [50].

##### 2. PI3K/Akt signaling pathway

Another critical pathway regulated by EGFR is the PI3K/ AKT pathways, which have been discovered to be among the most often misregulated signaling pathways in various human malignancies, including breast cancer [51]. Researchers have found that PI3Ks play essential roles in controlling numerous cellular processes necessary for cancer development, including metabolism, cell survival, proliferation, differentiation, and motility [51]. The PI3K-AKT is crucial for the control of intracellular physiological processes. This signaling pathway is stimulated by several oncogenes and growth factor receptors, including MET, KIT, EGFR, and ERBB [50].

##### 3. ERBB Signaling pathway

Activation of the ERBB family of receptors forms a significant group of related signaling pathways that control breast cancer’s proliferation, survival, angiogenesis, and metastasis. ERBB receptors modulate mammary gland growth and are often intensified, mutated, and overexpressed in breast cancer [50]. Since they have essential functions in regulating tumor proliferation, survival, and metastasis, the ERBB family could be a therapeutic target in breast cancer. Recent studies show that the ERBB family’s signaling irregularities and mutations are important in escaping antitumor immunity in the cell process [50].

##### 4. Estrogen receptor signaling

The majority of human breast cancers start as estrogen-dependent. The estrogen signaling and the estrogen receptor (ER) affect breast cancer progression [52]. Many studies reveal that ER signaling is intricate and involves extranuclear and coregulatory proteins. ER-coregulatory proteins are strictly regulated in a healthy state, with cancer being the primary reported instance of misexpression. Resistance to endocrine therapy results from the complexity of the control of estrogen signaling and the interaction of other oncogenic signaling pathways. Alternative strategies that target new molecular mechanisms are critical to overcoming this current and urgent gap in therapy [52].

##### 5. P53 signaling pathway

Mutation in p53 is one of the most common genetic changes detected in human neoplasia. P53 mutation in breast cancer is linked to more aggressive cases and lower overall survival. The tumor suppressor P53 recreates a crucial role in controlling cancer [53]. In luminal-type breast cancer, P53 is mutated infrequently but still functions normally. Posttranslational modifications like ubiquitination have a significant role in regulating the activity of wild-type P53. It has been demonstrated that several ubiquitin ligases manage the protein stability and ubiquitination of P53. These findings indicate direct P53 regulators and a possible therapeutic target for breast cancer [53].

##### 6. EGF signaling pathway

The function of VEGF signaling in breast cancer is not restricted to angiogenesis. This signaling pathway in breast carcinoma cells is vital for the ability of these cells to escape apoptosis and progress toward invasive and metastatic situations [53]. Cells capable of VEGF signaling has an advantage for survival and dissemination of breast cancer. These findings suggest that VEGF and VEGF receptor-based therapeutics could target tumor cells in addition to targeting angiogenesis [54].

##### 7. Wnt signaling pathway

Wnt signaling is vital in controlling embryonic organ development and cancer progression. Recent studies showed that Wnt signaling is involved mainly in breast cancer proliferation and metastasis processes [55]. Recent studies also revealed that Wnt signaling is vital in breast cancer therapeutic resistance, immune microenvironment regulation, stemness maintenance, and phenotype shaping. These accumulative pieces of evidence, which show the relationship between the Wnt signaling pathway and breast cancer, encourage researchers to use Wnt-targeted small molecules as attractive therapeutic targets in clinical trials for breast cancer treatment [55].

##### 8. MAPK signaling pathway

The MAPK pathway is a highly protected module that plays an essential role in different processes such as migration, cell proliferation, differentiation, and migration [51]. Any irregular activities in this pathway may cause many diseases, such as multiple types of cancer. The low frequency of Ras/MAPK pathway mutations in breast cancer indicates that the route is neither necessary nor utilized by breast carcinomas or the normal epithelial cell progenitors of breast cancers to initiate sustained growth and survival programs [56]. Instead, cells of mammary origin use alternative parallel or redundant signaling cascades to achieve the same results. For instance, alterations in the parallel and highly connected PI3K/Akt pathway are much more frequent in breast cancer but occur in different types of breast cancer [56].

##### 9. FOXO signaling pathway

Most breast cancers express the estrogen receptor, and endocrine-based therapies have remarkably improved patient outcomes. Endocrine resistance, however, is a very common occurrence [57]. Studying the pathways that control the hormone sensitivity of breast cancer cells is critical to improving the effectiveness of endocrine therapy. Recent studies showed that the PI3K/AKT/FOXO signaling pathway is a key player in the hormone-independent growth of many breast cancers. The PI3K/AKT pathway activation, as a driver of breast cancer growth, induces down-regulation of FOXO tumor suppressor functions [57].

After discussing and showing the connection between these pathways and breast cancer, we introduced the significant module among the 500 selected modules in this work with a p-value less than 10 * *E*^-1^ as the significant module related to breast cancer. Table 6 shows the set of these significant modules and their p-value, density value, the average score of the genes for each module, and the signaling pathways associated with each module.

**Table 6.**
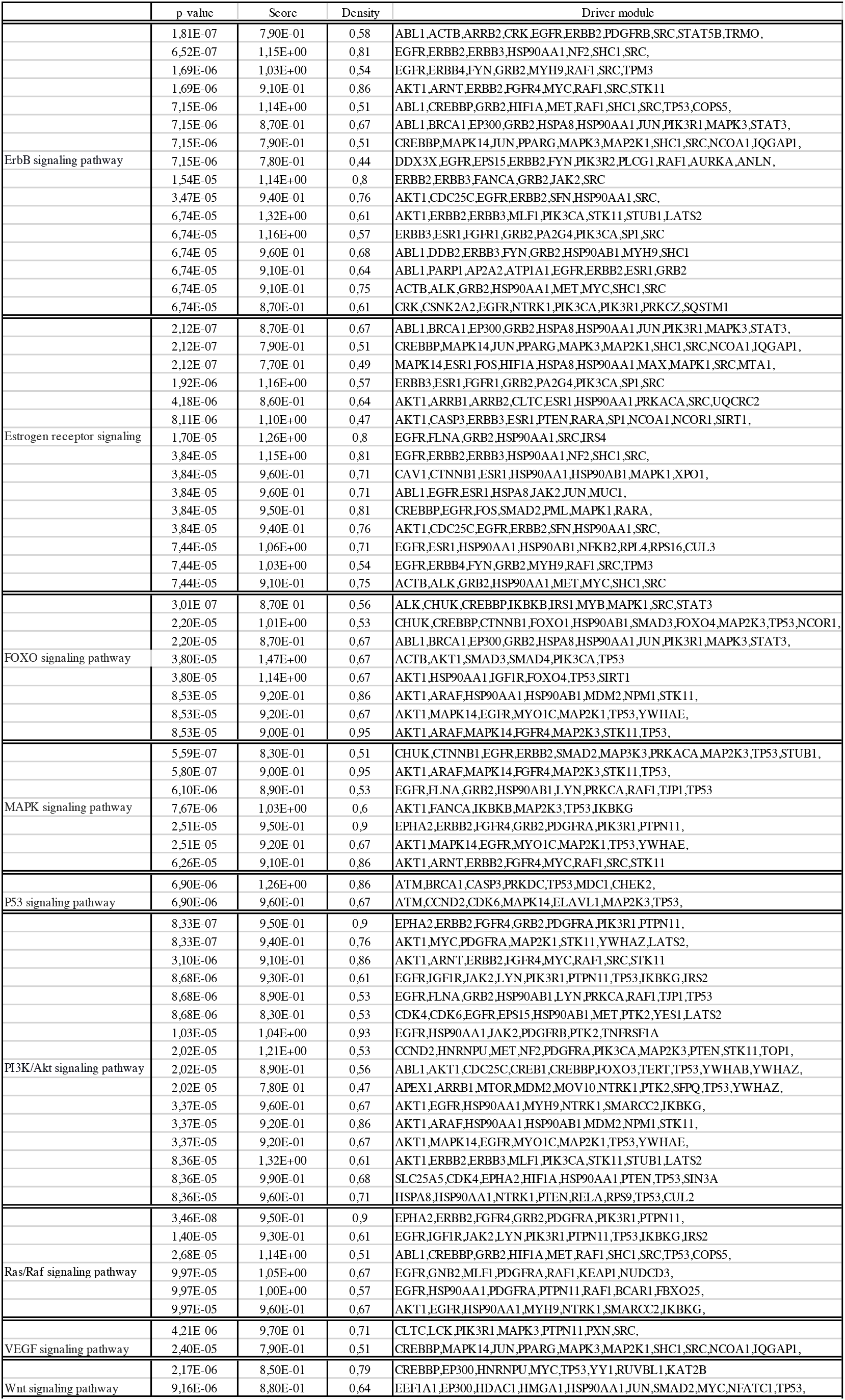
List of 75 driver modules, density value, score, and p-value for each module.

## 4. Conclusion

Breast cancer is the second cause of cancer death in developed countries and the most frequently diagnosed malignancy in women. While improvements in early detection and therapy alternatives have significantly reduced mortality, a patient’s reaction to treatment depends on the disease’s genetic variability. Recent mutation investigations demonstrated the presence of hundreds of low-frequency mutated genes, in addition to known driver genes. This finding encourages research into these genes’ effect on the disease’s pathogenesis.

In this work, we proposed two novel algorithms; in the first algorithm, we identified breast cancer driver genes with different mutation frequencies to find complete mutation patterns. In the second algorithm, we identified driver modules related to breast cancer. We used the set of mutated genes in breast cancer patients to find breast cancer driver genes. We built a biological network corresponding to the set of mutated genes, the network nodes corresponding to the set of mutated genes, and the network edges corresponding to the cases with common mutations in at least one case. Then, we defined topological and biological features for each node and proposed a novel machine-learning algorithm based on feature selection to assign scores to each gene. We also constructed a simulated data set to evaluate our proposed SPNFSR algorithm. The results of our investigations showed that the scoring method of SPNFR was not random. The study of high-score mutated genes showed that our selected gene set includes both high and low-frequency mutated genes. We also checked 30 genes with high scores and low frequency identified by our algorithm. Then we introduced the breast cancer network with respect to the biological process corresponding to the high-score genes and identified a set of 262 mutated genes in the breast cancer network. To better evaluate our algorithm, we studied and evaluated the obtained candidate driver gene set from different aspects. We have provided a comprehensive comparison and a complete review of this set of genes and have shown that this set of genes has a high agreement with the driver genes of benchmark sets. We also evaluated the performance of different algorithms with different approaches and their results on these benchmark sets (Table 5). Our investigations showed that our algorithm identified 8 genes as breast cancer driver genes that most algorithms do not confirm. We studied the association of these genes with breast cancer separately and suggested them for further investigation in clinical studies.

The other goal in this study is to identify the modules associated with breast cancer. Since the breast cancer network is dense, the module identification algorithms could not perform well on this network. Therefore, we have proposed a Monte Carlo randomization algorithm to identify high-density networks with high scores in the breast cancer network. We found 500 high-density networks with high scores. For more investigation, we calculated the p-value of each of these obtained modules according to the well-known pathways identified in cancer. We presented 75 significant modules with high coverage in cancer pathways in Table 6. Identifying the driver genes and modules of breast cancer will provide a complete insight into the cause of breast cancer and its development.

